# Infinite storage capacity of memory-like mechanisms in the hippocampus: a cryptographic perspective on animal’s caching and retrieval behaviors

**DOI:** 10.1101/2021.08.11.455910

**Authors:** Sharon Mordechay, Oren Forkosh

## Abstract

The brain’s extraordinary abilities are often associated with its ability to learn and adapt. But memory and plasticity have their limitations, especially when faced with tasks such as retrieving thousands of food items such as in the case of scatter-hoarding animals. Here, we suggest a brain mechanism that works by utilizing cryptographic principles in lieu of plasticity. Rather than memorizing the locations of their caches, as previously suggested, we propose that cache-hoarding animals use a single cryptographic-like mechanism for both caching and retrieval. The mathematical model we developed functions similarly to hippocampal spatial cells, which respond to an animal’s positional attention. We know that the region that activates each spatial cell remains consistent across subsequent visits to the same area but not between areas. This remapping, combined with the uniqueness of cognitive maps, produces a persistent crypto-hash function for both food caching and retrieval. We show that our model is consistent with previous observations, such as animals’ ability to prioritize food items that are perishable or by their nutritional value. The model makes several measurable predictions regarding scattered hoarding and what factors can limit an animal’s retrieval success. Finally, while focusing here on scatter-hoarding, the mechanism we present might be utilized by the brain in other ways providing essentially infinite retention capacity for structured data.

## Introduction

Animals have much to hide in order to survive. Some species evade potential predators or prey by finding cover or by using camouflage (1), mimicry, and other means of disguise (2). Others conceal their eggs or offspring, mask an illness or an injury to avoid being targeted by predators (3), or stash valuable resources, such as food.

Scatter hoarding is probably the largest-scale manifestation of secretive behavior in the animal kingdom. Many species of animals engage in this behavior, which involves storing food at multiple cache sites to preserve it for times when food is scarce (4). Several bird species, such as the Siberian tit (*Poecile cinctus*), were claimed to cache over 500,000 items per individual in one year (4). While much of the research on scatter-hoarding was conducted on birds, this behavior is not specific to them (4): squirrels (5), chipmunks (6), and even foxes (7,8) stash food for times of need. As caching sites cannot be defended, the success of this strategy is often contingent on an animal’s ability to keep the stashes away from prying eyes and hard-to-find (5).

The ability to retrieve items from cache sites depends on spatial information such as visual cues. In (9), black-capped chickadees (*Poecile atricapillus*) were placed in an enclosure and their food-caching behavior was tracked. Object rearrangement around the enclosure greatly impaired the chickadees’ ability to find their cache sites; manipulation of prominent global landmarks (large cardboard cutouts and a poster) had a much stronger effect on the birds’ retrieval performance than small proximal objects (5 cm squares). Shifting objects by as little as 20 cm to the right significantly decreased the chickadees’ ability to recover the food. Moreover, in almost 70% of the cases, the birds searched within 5 cm of the location implied by the more prominent landmarks, with a mean displacement of around 20 cm. Assuming this is approximately the caching resolution, the finding indicates that a small area of 10}10 meters can hold as many as 2,500 potential caching sites.

The hippocampus plays a central part in the remarkable cognitive feat of caching (10). This is not surprising, as the hippocampus is known to be involved in processing spatial information in the brain (11). A large subpopulation of neurons within the hippocampus in animals such as mice, rats, and bats exhibit *place-cell* behavior; that is, they increase in their spike rate in response to the animal’s entering a specific region within a given site (usually, but not always, one region per cell). The region activating each place cell often changes when the animal moves to a new area, often in an unpredictable manner. However, if the animal returns to a site previously visited, the place cell’s receptive fields also return to their previous arrangement, and this change happens practically instantaneously. This remapping of the receptive field within a given environment is mostly insensitive to landmark manipulations. In primates, we usually find a related type of cells referred to as spatial view cells. These cells respond remotely when an animal is gazing at a specific region, independently of the animal’s location or head direction (12).

A well-known homolog to the mammalian hippocampus also exists in birds, with similar involvement in spatial and episodic memory (13). Hippocampus size in birds was found to correlate with birds’ ability to stash food. Although the interpretation of this correlation is under debate (9,14), animals that used more cache sites generally had a larger hippocampus than non-caching bird species (15,16). In addition, even within the same species, the size of the hippocampus was found to be larger in individuals dwelling in harsher environments that makes them more dependent of the cached food (17). Moreover, hippocampal neurogenesis has a seasonal element and seems to correlate with caching activity throughout the year (18). For many years, the spatially responsive cells found in avian brains were less related to a fixed position in space and more related to the challenge the animal faced such as the position of a goal within a maze (19). Only very recently the existence of place cells was demonstrated in the tufted titmouse (*Baeolophus bicolor*) (20).

Taken together with the fact that the hippocampus is involved in memory, these observations have led researchers to hypothesize that caching requires some form of spatial and episodic memory (21).

Yet as birds and other animals need an internal mechanism to guide them to stash food in specific locations, the same mechanism can also be used to direct them to the exact same locations while retrieving the food in that area. A mechanism, or mapping, that can facilitate efficient hiding and retrieval of multiple cache sites without relying on memory would need to have several basic properties. From a theoretical perspective, the class of methods that achieve this is known as cryptographic-hashing functions (or crypto-hashes, for short), which, as the name suggests, are comprised of two components: hashing functions and cryptographic keys (22).

In this context, hashing refers to a class of functions that map arbitrarily complex data (images, texts, audio files, and others) to a fixed size lower-dimensional representation. Computer applications often use hashes to store objects in memory efficiently by mapping them directly to a memory location (a type of mnemonic device). In the case of food caching, hashing can be used to map a set of landmarks within and around an area onto a selected caching site within that area (Figure 1A). Efficient hash functions are such that the probability of assigning different cues to the same output is kept to a minimum; This property reduces the possibility of collisions and redundancies that can occur when two different inputs result in the same output. It also makes better use of all the available resources – in the case of animals allowing the use of the entire area for caching.

**Figure 1.**
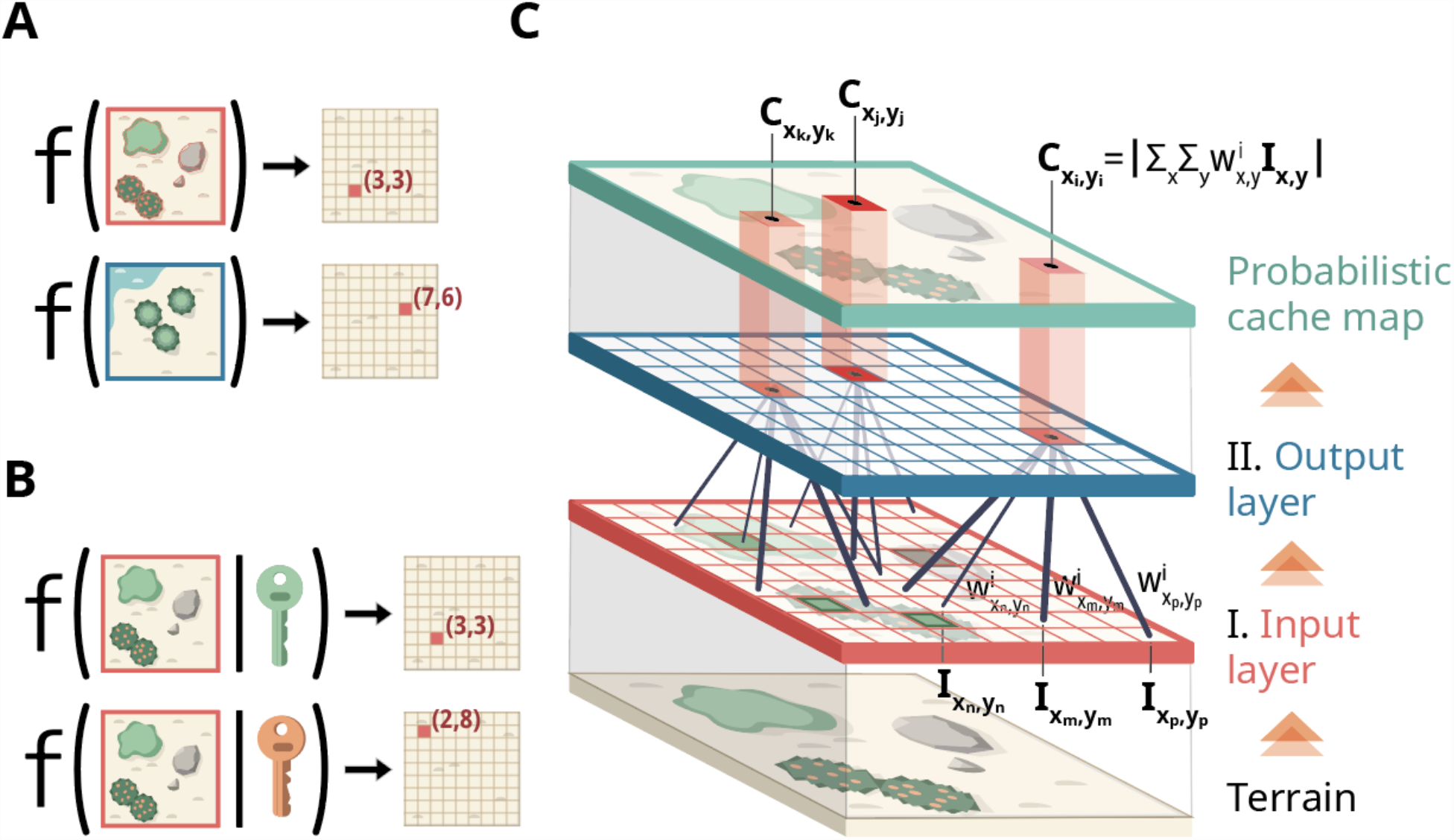
Cryptographic mechanisms in the brain. (A) Hash functions take high-dimensional objects (a map of a complex terrain, for example) and map them into a low-dimensional representation such as a point within that terrain. (B) Crypto-hash functions also include a private key that makes mapping the same object unique across individuals with different keys. (C) Crypto-hashing in a two-layer neural network. Each neuron *I*_*x, y*_ in the input layer represents landmarks within a small square area. We assume two types of landmarks, which we refer to as ‘trees’ and ‘rocks’. Neurons that point to trees are assigned a positive value, rocks get negative values, and if no object is within the neuron’s receptive field, it is set to zero. The absolute value of *I*_*x, y*_ corresponds to an object’s prominence; thus, prominent trees will get +1, smaller trees – 0.5, and small rocks may be assigned a value of -0.3. The spatial output layer is a 2D mesh that assigns a caching score to each location within the site. The higher the score, the more likely this location would be used for caching. Each neuron in the output layer is innervated by a small number of input neurons. In all of our simulations, we matched this number to the number of landmarks the model uses.

Unlike typical hash functions, crypto-hashes incorporate an additional factor – a private key - that renders the mapping unique to the key owner (22). Assuming no two individuals have the same key, it also means that the mapping will result in unique caching sites within the same area (Figure 1B). Another valuable property of crypto-hashes is that it is often difficult to infer the key from a small number of examples, so even if another animal finds several caches, it will not be able to deduce the location of all others.

A brain mechanism such as this may serve as a mnemonic device (as suggested in (23)) or possibly replace the need for memory altogether. Such a pseudo-random approach is much simpler than remembering hundreds of thousands of stashing sites while still supporting all the existing empirical evidence. The guidance is based on prominent landmarks in the terrain, such as trees and rocks, which are not likely to substantially change over time, and can be used in the subsequent cache retrieval. And we already know of a specific set of neurons to be able to do precisely this – the previously mentioned hippocampal spatial cells. Spatial cells are unique to each individual, they assign scores and rankings (using spike rates) to different locations within each area, they persist over time, and remap when in the same area. We show how these properties allow animals to find their cache sites efficiently and secretly.

## Results

### Biological crypto-hashing network

A straightforward and biologically plausible realization of crypto-hashing is through a sparsely connected two-layer neural network (Figure 1C; see supplementary information). The first layer, the input layer, represents the visual cues (landmarks) within a given patch of land (Figure 2A, 2B). The second layer is a 2D lattice of spatial neurons, in which each neuron points to a specific location in a given area (Figure 2C). The firing rate of each spatial neuron corresponds to the likelihood of choosing its particular location as a caching site. The spatial neurons are sparsely innervated by the input neurons.

**Figure 2.**
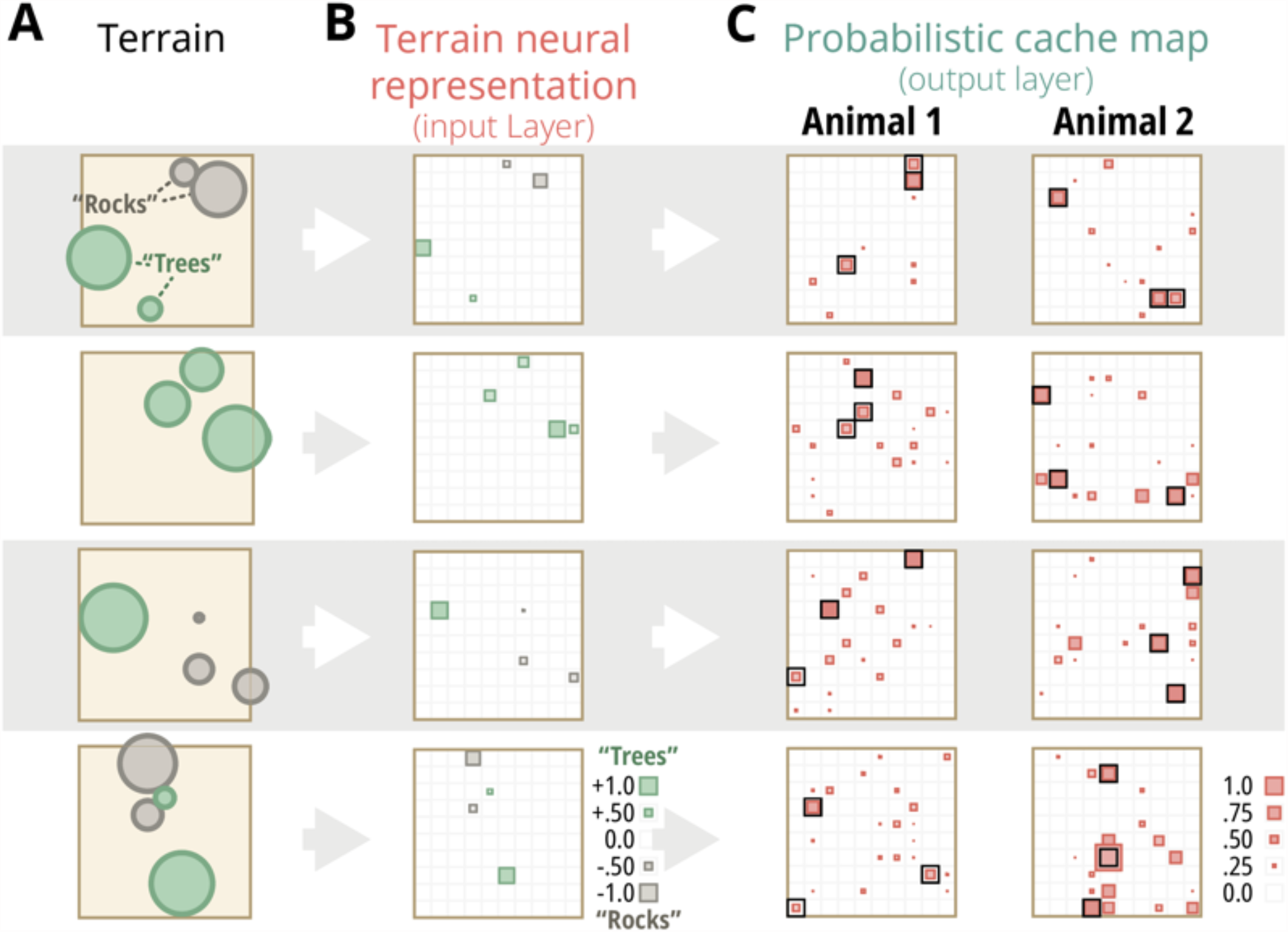
Using spatial features to map cache sites without requiring brain plasticity. (A-C) Four examples of the outcome of a crypto-hash neuronal network (in rows). (A) We simulated a terrain with four prominent landmarks by randomly choosing four cells within a 10}10 grid. The cells were assigned random values between -1 and +1, so that the absolute value represents the prominence of the spatial feature (cells with values close to +1 and -1 being the most prominent), and the sign represents the type of object. We refer to positive-valued cells as “trees” and negative cells as “rocks” for brevity. (B) The representation of the landscapes from (A) in the neural network’s input layer. The size of the colored inlaid boxes represents the object’s prominence and their color its sign (green for positive values or “trees”, and gray for negatives or “rocks”). (C) The output of the target layer of two randomly chosen neural networks (Animal 1 and 2) in response to the inputs in (B). The output layer creates a unique probabilistic map of possible cache sites.

In our model, we set the number of connections to a constant, typically equivalent to the number of landmarks the model uses (usually four). This sparse connectivity helps maintain a low number of potential cache sites by the output layer. A simple equation can summarize the activity of each spatial neuron

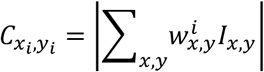

where 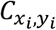 is the score, or spike-rate, of the spatial neuron pointing to coordinate *x*_*i*_, *y*_*i*_ of the output grid; *I*_*x,y*_ is the input from coordinates *x, y*; and 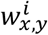 represents the strength of the connectivity between *I*_*x, y*_ and 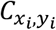. The strength, or weight 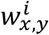, of the connections was assigned randomly at between one and minus-one. The inputs were also set in the range one and minus-one, where the absolute magnitude represents the landmark’s prominence (one is very prominent and zero is designated as not noticeable). The sign represents the landmark type, for example positive values represent “trees”, and negatives represent “rocks”. The scores of the output neurons 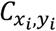 effectively determine the probability of their target area being used as a cache site. We use the absolute value to keep the outputs positive, although it has no computational benefit to the model. Nor is the choice to distinguish between two types of objects by allowing negative inputs.

This neural network is a crypto-hash function, as it fulfills the three essential properties: (1) It maps a complex terrain into a point with minimal overlapping probability across the terrain or (2) across subjects (Figure 3A, 3B), and (3) reconstructing or decrypting the mapping from examples is difficult. The third point stems from the fact that the connection to each output-layer neuron is chosen randomly and independently of the others. Thus, it is effectively equivalent to holding a unique key for each neuron.

**Figure 3.**
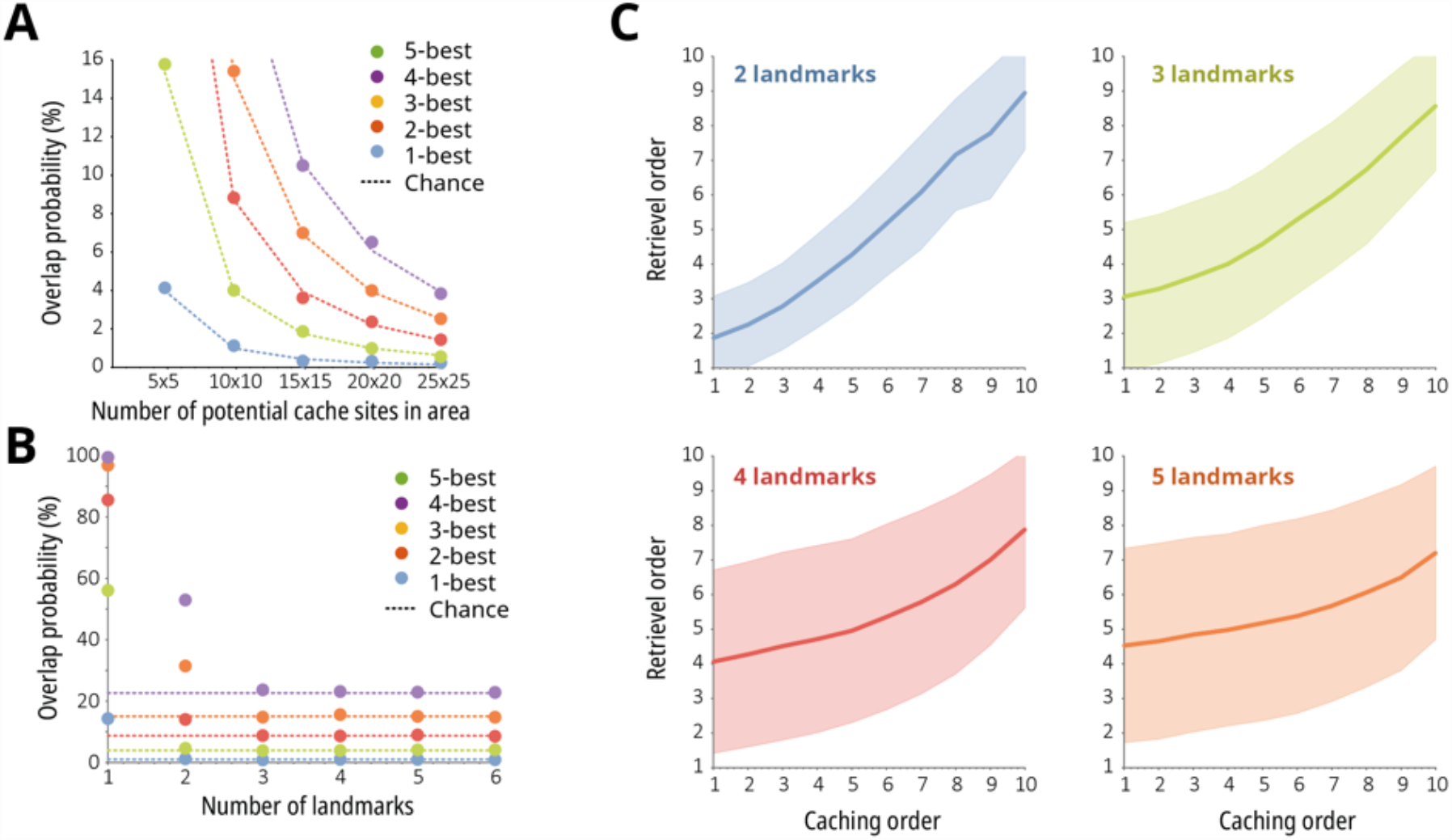
Properties of a crypto-hash neural network. The probability of overlapping sites between two random networks as a function of (A) potential cache sites and (B) the number of landmarks the network uses. The different colors correspond to the number of caches used in each area. The dotted lines are the overlap as expected from a random choice of sites. (C) *The probabilistic nature of the target layer scores allows the network to maintain temporal dynamics*. We assume that caching and retrieval order are determined by the target-layer score, from highest to lowest. If the choice is not absolute but probabilistic, we still get the same temporal dynamic in both phases. The shaded area around each line represents the standard deviation.

### Benefits of probabilistic maps

The result can serve as a traditional crypto-hash function by choosing the target neuron with the highest score as the mapping outcome (see Materials and Methods). However, a probabilistic map with multiple outcomes of varying probabilities – apart from being more biologically feasible – also provides several benefits. The most straightforward benefit is that it allows for an arbitrary number of caching sites within each area by choosing the spatial neurons with the top scores.

Another valuable property of probabilistic maps is that they allow the addition of temporal considerations into caching behavior (21). Assuming the order of food recovery starts with locations that have higher scores, items with higher nutritional values or perishable items (such as dead insects, as opposed to seeds) could be stashed in places with higher scores – making them more likely to be recovered prior to items assigned to lower-scored locations (Figure 3C). In addition, avoiding previously excavated sites requires only memorizing the score of the last excavated and choosing only sites with a lower score (we refer to this behavior as bookmarking).

Finally, caching maps should be allocentric and invariant to the animal’s position. A straightforward approach to achieving such invariance was suggested in an elegant paper about geometric hashing (24). The method there is based on choosing two prominent objects in the area and using them to scale and align all landmarks. The vector connecting the two most prominent landmarks defines the direction axis and the distance between them sets the scale unit. Using this approach, we can obtain a model that is insensitive to affine transformations.

## METHODS

### Landmarks and terrain

While the algorithm is not sensitive to the number of landmarks, for the sake of simplicity, we assumed a fixed number of landmarks within each area - four, in this paper. These landmarks were divided into two categories, which we refer to as “trees” and “rocks”. The locations of the landmarks were chosen randomly and uniformly from an n-by-n square grid (in most cases we use n=10). Each landmark was then assigned a random value between minus one and one that signifies the objects category and prominence: Cells that have positive values are referred to as trees (values between zero and one), while rocks had negative numbers (between zero and minus one). The absolute value of each landmark signifies its prominence - so prominent trees have values closer to one and prominent rocks have values closer to minus one. Zero marks a no-object. The outcome is a sparse n-by-n matrix *S*_*n*×*n*_ with values that vary between minus one and one.

### Crypto-Hash Functions

Hash functions map data with arbitrary dimensions to a fixed-length value (22). In mathematical terms, a hash function *g*_*p*_(*<u>s</u>*) is such that

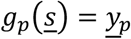

where *<u>s</u>* ∈ *S* is a vector of arbitrary length, and *<u>y</u>*_*p*_ = (*y*_1_, *y*_2_, …, *y*_*p*_) ∈ *Y*_*p*_ is a vector of a fixed-length *p*. Since the length of *<u>s</u>* is often larger than that of *<u>y</u>*_*p*_ hash-functions can be viewed as a special case of dimensionality-reduction.

An optimal hash function is such that the probability of mapping two inputs onto the same output is minimal, or, equivalently, that all outputs values should have (roughly) the same probability. This principal of uniformity can be formulated as

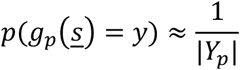

where |*Y*_*p*_| is the cardinality (number of elements or size) of the set *Y*_*p*_ of all possible output values. Because of their uniformity, hash functions are often used in data storage and retrieval tasks as they allow data access at nearly a constant time while requiring a storage size that is only slightly larger than the space needed to store the data itself.

Crypto-hash functions introduce an additional term, a private-key *k*, to the basic hash function *g*_*p*_ (*<u>s</u>*; *k*) = *<u>y</u>*_*p*_. The key ensures that the mapping is unique, i.e. the probability that the same inputs produce the same outputs for different keys is close to chance. Crypto-hash functions, like hash functions in general, are deterministic, meaning that the same combination of input and key will always produce the same output value. However, crypto-hash mappings are also one-way-functions, meaning that they are difficult to invert; knowing an output value gives very little information about the input or key.

### Crypto-Hash Neural Network

Choice and retrieval of cache sites is based on prominent landmarks within a terrain. Assuming *S*_*n*×*n*_ is the representation of the current area’s terrain (see the ‘simulated terrain’ section), our crypto-hash function can be defined as

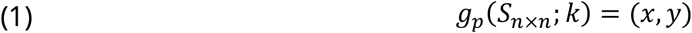

where (*x, y*) is the cache coordinate within the area so that *x, y* ∈ {1, …, *n*}, and *k* is the crypto-key.

A straightforward and biologically plausible to achieve this is using a neural network. We define a two-layer network where, for simplicity, the neurons on both layers are organized as a grid with *x, y* indices. The value of each neuron in the first layer *I*_*x, y*_, which is the input layer, is set according to the corresponding area tile or the (*x, y*)’th cell in *S*_*n*×*n*_. Each output layer neuron 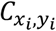 was sparsely connected to the input layer, and the weights 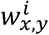 were randomly distributed between minus-one and one. The value of output neurons is the absolute value of the weighted sum of their inputs

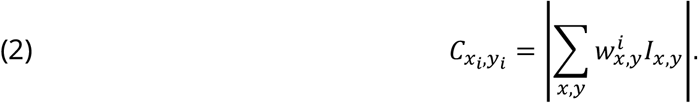

In order to get a crypto-hash function in the form (1) we can take the index of the maximal value or 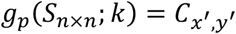.

In our case, a unique key *k* is obtained from the random choice of weights between the neural network’s layers.

However, as we mentioned in the text, keeping the function as a probabilistic mapping like in equation (2) has several benefits and this is the form we use in the paper.

## DISCUSSION

Although spatial cells, such as place cells have been extensively studied, the mechanism we presented is one of the few explanations for how the brain might utilize these cells. In this study, we propose a novel perspective suggesting that spatial cells function as the brain’s crypto-hash functions, facilitating animals to conceal food in distinct cache sites and later retrieve it. So far, there has been no theory that explains what principles guide animals when choosing cache sites; If it was merely a question of optimality, all animals might end up choosing the same sites, which would lead to theft (or klepto-parasitism). We have demonstrated how our model can explain previously observed complex behaviors related to scattered hoarding, such as retrieving perishable food items first.

While our focus here is on scattered hoarding, a similar mechanism might also allow animals of the same species to find each other in complex environments. Simple crypto-hash functions that are conserved across individuals of a species can help them navigate to the same place when looking for mates or other social activities. The same function-driven mechanism (rather than memory-driven) may also help steer migratory animals. Finally, since the hippocampus is involved in abstract knowledge in addition to spatial information (11), the scope of decision-making might be much broader; Since diversity is a key characteristic of all living system, it is tempting to think that humans’ individualistic tendencies might also be somehow related to our proposed brain circuitry.

While the work we presented is theoretical, it raises some obvious predictions. First, knowing the spiking patterns of spatial cells will enable us to determine cache site locations. Moreover, if we know the remapping between sites well, we can use it to decrypt the internal circuitry and predict cache sites in a new site that the animal is yet to visit. Finally, we predict that the location of cache sites within a given area would be consistent across multiple hide and retrieval iterations.

The instinct to choose cache sites that are both unique and obscure has a clear evolutionary advantage. We therefore fondly suggest addressing this movement pattern as *cryptotaxis* and the neurons involved as *crypto-cells*.

## Funding

This work was supported by the Israeli Science Foundation (ISF; Grant 2505/20) and the Ring Center for Interdisciplinary Environmental Research.

## Notes

### Competing Interest Statement

The authors have declared no competing interest.

### Summary of Updates

Added analysis and made several corrections to math and figures

